# Kaposi Sarcoma herpesvirus reactivation and replication are dispensable for cell-lineage specific tumorigenesis in whole genome transgenic mice

**DOI:** 10.64898/2026.06.10.731437

**Authors:** Kyle W. Shifflett, Anthony B. Eason, Huanjuan Su, Johann W. Schneider, Yongbaek Kim, Maria C. White, Zhigang Zhang, Blossom Damania, Dirk P. Dittmer

## Abstract

Herpesvirus reactivation and replication depend on an ordered cascade of gene expression. The viral immediate-early replication and transcription activator protein (Rta) is necessary and sufficient to initiate this process. For Kaposi sarcoma herpesvirus (KSHV), Rta-interacting and Rta-regulated proteins are thought to contribute to cellular transformation. Using a novel, whole-genome transgenic mouse model, we show that KSHV tumorigenesis is independent of Rta *in vivo*. Rta-deleted, KSHV bacmid transgenic (tgKSHVΔRta) mice developed latency-associated nuclear antigen (LANA)-positive vascular sarcomas with the same endothelial cell-lineage composition and transcriptional profile as human Kaposi sarcoma (KS). The viral miRNAs were expressed in the tumors, as were all canonical latency and the K1 and K15 oncogenes. Importantly, this system is the first to model both KS and lymphoma, reflecting the development of both malignancies observed in KSHV-infected individuals. These results separate virus replication from KS tumorigenesis, which can proceed independently so long as the virus genome is maintained in a susceptible cell.

**Significance:** Kaposi sarcoma (KS) is the most common cancer in people living with HIV worldwide. Like all herpesviruses, KS-associated herpesvirus (KSHV) has a latent and lytic phase. Hitherto, viral lytic genes are thought to play a role in KS. The KSHV replication and transcription activator (Rta) is both necessary and sufficient for reactivation and lytic viral replication, but Rta was not required for vascular tumor development in a whole virus genome mouse model. This new tgKSHVΔRTA mouse model separates virus replication from viral tumorigenesis and defines a minimal set of viral oncogenes. This model enables mechanistic studies *in vivo*, and treatment modalities to be evaluated within a complete immune and stromal microenvironment.

## Introduction

Kaposi sarcoma (KS) defined the acquired immune deficiency syndrome (AIDS) epidemic in the US and Western Europe ^1^. In KS-associated herpesvirus (KSHV) endemic regions of Sub-Saharan Africa, KS manifests independently of HIV infection. Here, in children, KS is localized to lymph nodes, and skin lesions are often absent ^2,3^. In “classic KS” the incidence increases exponentially with age, absent overt immunosuppression. Irrespective of natural history, all forms of KS share a common histopathology. KS is a tumor of endothelial cells (EC) ^4–7^. KS lesions span a spectrum of microenvironments: some resemble a loose assembly of disorganized ECs (patch and plaque type), whereas others are composed of dense nodules of LANA-positive, so-called KS spindle cells ^8–10^. All KS lesions contain the KSHV genome and express the latency-associated nuclear antigen (LANA) ^11^; however, no mutational signature has been identified in KS ^12^, suggesting that the viral genome is necessary and sufficient for tumorigenesis.

Whether viral replication is required for cellular transformation or merely a means to infect people and establish molecular latency in susceptible cell lineages has remained an open question since the discovery of KSHV. Demonstrating KS development in the absence of viral replication or viral late protein expression would better define KSHV tumorigenesis, thereby narrowing the field of possible KSHV oncogenes as potential targets for therapeutic interventions.

The KSHV replication and transcription activator (Rta) is necessary and sufficient to initiate viral DNA replication and the release of infectious particles ^13,14^. Rta binds to consensus sequences in the viral genome either directly or via Recombination Signal Binding Protein for Immunoglobulin Kappa J Region (RBPJκ) ^15,16^. RBPJκ allows Rta to bind both cellular and KSHV promoters. To test the hypothesis that viral DNA replication, Rta itself, or Rta-dependent genes are required for tumorigenesis, we generated Rta-deficient whole-genome KSHV-transgenic mice (tgKSHVΔRta). Like our first-generation KSHV transgenic mice ^17^, the tgKSHVΔRta mice reproducibly develop a sarcoma with endothelial differentiation that, by molecular analysis and histopathology, is indistinguishable from human KS.

## Results

### KSHV whole genome, Rta-deleted transgenic mice develop murine KS

Rta exon 2 was deleted from a wild-type KSHV bacmid ^18^ to yield BAC16ΔRta (Figure 1A, B). In HEK293 cells, BAC16ΔRta expressed LANA but did not express the lytic protein K8α, unlike WT BAC16 DNA, which expresses both latent and lytic proteins (Figure 1C). In the iSLK-RTA cell line, which carries Rta in trans, both wild-type BAC16 and BAC16ΔRta expressed similar levels of K8α (Figure 1D). Microinjection of BAC16ΔRta into FVB/NJ zygotes yielded nine founder animals (Supplemental Table 1), three of which (33%) had offspring. Two founders (22%) developed angiosarcomas and pleural effusions due to nonspecific weakening and dilation of blood and lymphatic vessels, known as angiectasis, phenotypes previously observed in tgKSHVwt and tgKSHVΔmiR mice ^17^. One line, tgKSHVΔRta, stably transmitted the transgene and developed tumors with ∼18% incidence in the first generation. Most tumors were localized to the head and neck region, and edema was common (Supplemental Table 2). PCR confirmed the genotypes (Figure 1E). Whole-genome sequencing confirmed the absence of second-site non-synonymous mutations and the preservation of multiple copies of terminal repeat sequences, as well as the deletion of Rta (Figure 1F). The transgene integrated into chromosome 11 and did not disrupt an essential locus (Figure 1G).

**Figure 1:**
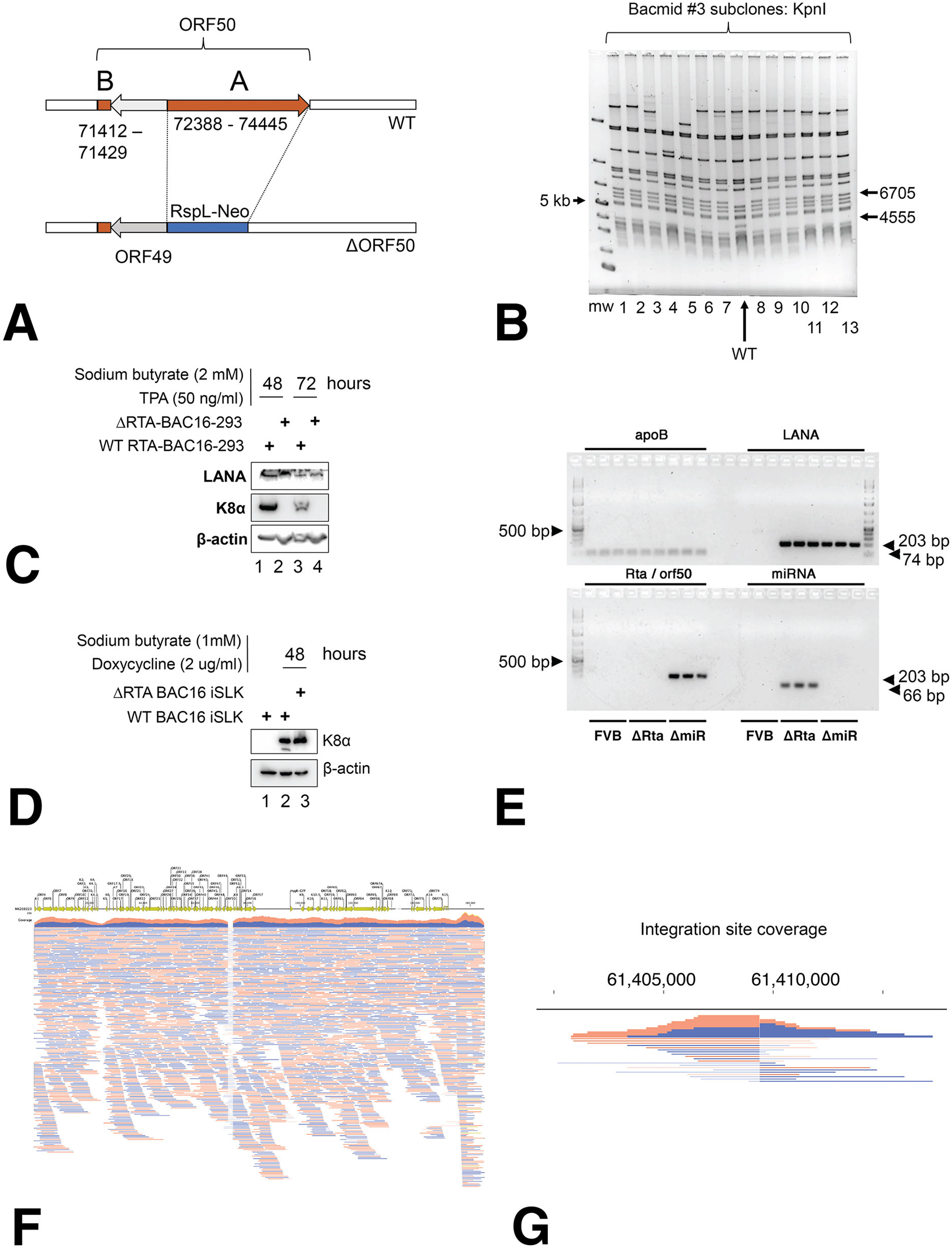
Construction. (A) The recombination strategy and (B) KpnI restriction digest of BAC16ΔRta bacmid #3. Arrows show bands of 6,705 bp (gained) and 4,555 bp (lost). (C) Western blot of HEK293 cells carrying BAC16 or BAC16ΔRta, after induction with sodium butyrate and TPA. (D) Western blot of iSLK carrying BAC16 or BAC16ΔRta, with (1) no induction or (2,3) 48 hours after RTA induction. (E) PCR of KSHV genes (LANA, ORF50, miRNA) and a control (apoB) from transgenic and transgene-negative FVB mouse DNA. (F) Read coverage of the KSHV genome from transgenic mouse DNA. (G) Reads that map to both KSHV and mouse.

Histologically, KS is characterized by proliferation of spindle-shaped cells in fascicles, neovascularization, vascular leakiness, EC dysfunction, vessel dilation, and LANA staining. Figure 2A & B show a human KS biopsy with hematoxylin and eosin (H&E) and LANA staining. The tumors in the tgKSHVΔRta mice shared the same characteristics and are henceforth referred to as murine KS. Multiple dark, blood-filled vascular tumors were evident in subcutaneous and visceral locations. H&E staining showed densely cellular, spindle-shaped cells forming irregular vascular spaces filled with erythrocytes (Figure 2C). Tumor cells were LANA positive (Figure 2D). In human KS, LANA shows multiple nuclear speckles, or “LANA dots,” each indicating the terminal repeat (TR) region of an extrachromosomal episome (Figure 2E)^19,20^. In murine KS, only one to two “LANA dots” were visible within the nuclei of the murine tumors (Figure 2F), consistent with a single integration site for the viral TR. The elongated tumor spindle cells (Figure 2G) stained positive for LANA (Figure 2H, L). The cells stained positive for VEGFR-3 and LYVE-1 (Figure 2I, J), consistent with lymphatic EC (LEC) origin. Proliferation, as measured by Ki-67 staining, was low (Figure 2K).

**Figure 2:**
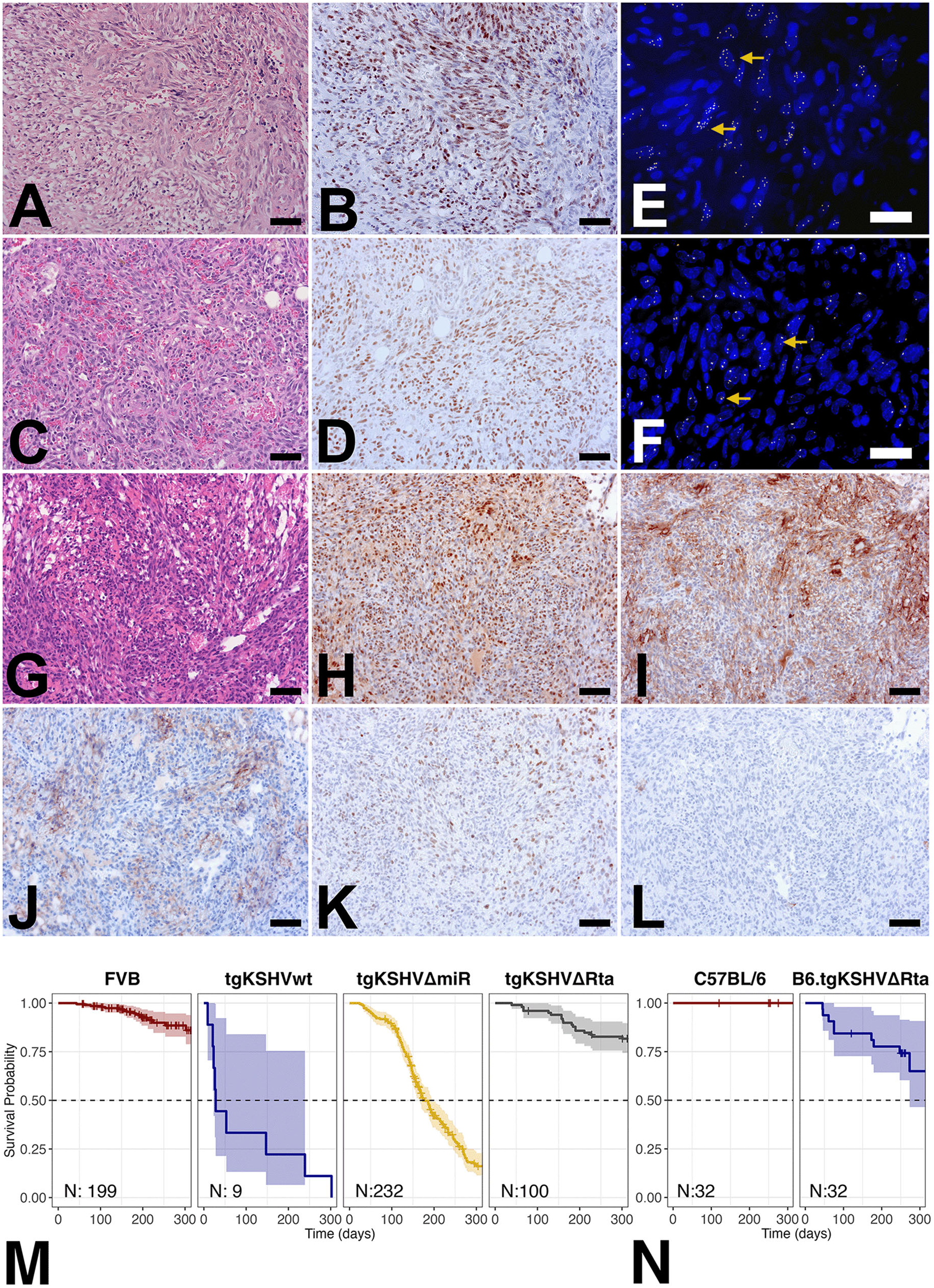
Histology. (A) H&E staining and (B) LANA staining of a human KS. (C) H&E and (D) LANA staining of murine KS (animal #506). (E) LANA IFA of human KS and (F) murine KS, yellow arrows point to LANA+ nuclei. (G) H&E, (H) LANA, (I) VEGFR-3, (J) LYVE-1, (K) Ki-67, and (L) no primary antibody staining on murine KS (animal #535) (black bars = 50 µm, white bars = 20 µm). (M) Kaplan-Meier plot of OS for transgene-negative FVB, tgKSHVwt F0, tgKSHVΔmiR, tgKSHVΔRta and (N) transgene-negative C57BL/6 and B6.tgKSHVΔRta.

KS has a variety of clinical and histological presentations, with lesions being classified based on location and distinguishing histological features ^21,22^. Applying these classifications to tgKSHVΔRta tumors, three major presentations could be discerned. (i) Liquid-filled tumors that ruptured upon puncture, which were defined by cavernous blood or lymph-filled spaces lined by a single layer of endothelial cells, reminiscent of lymphangioma-like KS ^23^ (Supplementary Figure 1A-F). (ii) Solid tumors in lymph nodes, reminiscent of pediatric or lymphadenopathic KS (Supplementary Figure 1G-L). (iii) Solid nodular tumors under the skin and attached to proximal tissues, including muscle, which were reminiscent of advanced, adult AIDS-KS (Supplementary Figure 1M-R). These presentations were not mutually exclusive; many animals had all three concurrently. 31 incident tumors from 17 mice across multiple generations were evaluated and graded by a human pathologist and a veterinary pathologist. 14 of 31 total tumors (48%) contained recognizable lymphoid tissue. Salivary gland tissue was diagnosed in 15 of 31 tumors (48.39%), typically at the periphery. The most common immune cells across all tumors [31/31 (100%)], based on H&E stain were small lymphocytes. Plasma cells were found in 27/ 31 (87%) of tumors (Supplemental Table 2). In lymph node-localized tumors, dense groups of lymphocytes were pushed to the periphery of the tumor due to the proliferation of VEGFR-3-positive spindle cells (Supplemental Figure 2A, B, E, F). In many such cases, Ki-67 staining showed increased lymphocyte proliferation at the tumor-lymph node interface (Supplemental Figure 2C, D) and in germinal center-like structures (Supplemental Figure 2G, H), indicative of activated lymphocytes. Expanded germinal centers appeared in lymphoid tissue within tumors (Supplemental Figure 2I, M, J, N) and in lymph nodes contralateral to tumors (Supplemental Figure 2K, O). Distal lymph nodes occasionally showed dilated vessels and extravascular erythrocytes (Supplemental Figure 2L, P), indicative of EC malfunction.

Tumor incidence remained stable across multiple generations (n=117), with 13% of animals developing tumors, 67% of which were female, a median of 2 tumors per mouse, and an average latency of 206 days (95% CI: 170…242, n = 15). Fatal angiectasis developed in two animals. Overall survival (OS) of tgKSHVΔRta (median of 390 days, 95%CI: 389–NA, n = 100) was lower than that of wild-type FVB (median survival was not reached, n = 199) (Figure 2M). This contrasted with tgKSHVΔmiR mice (median of 177 days, 95%CI: 166 – 196, n = 232) and tgKSHVwt founder mice (median of 28 days, 95%CI: 23 – NA, n = 9). In tgKSHVΔmiR mice, OS was driven by angiectasis, while in tgKSHVΔRta mice, survival was determined by neoplasia.

To test the hypothesis that the KSHV-dependent tumor phenotype is strain-invariant, the B6.tgKSHVΔRta strain was developed, which carries the KSHV transgene in the C57BL/6 background. These also developed LANA-positive murine KS. The difference in OS between the B6.tgKSHVΔRta mice and non-transgenic C57BL/6 littermates was significant at p≤0.001 (Figure 2N). 11/32 (34%) animals died of tumors by 300 days, while all non-transgenic littermates (n = 32) remained alive and symptom-free. This demonstrates that the KSHV-induced tumor phenotype is strain-independent.

### Transcriptional profiles of murine KS

13 tumors from tgKSHVΔRta were subjected to RNA sequencing and compared to 5 tumors from tgKSHVΔmiR mice. The tumors that developed in the two mouse lines differed across many transcripts (Figure 3A) and clustered separately (Figure 3B), except for tgKSHVΔmiR mouse #338, which was more similar to tgKSHVΔRta tumors. We evaluated mRNA levels for the LEC lineage markers CD34, Flt4/VEGFR-3, Lyve1, PDGFR-α (Pdgfra), Podoplanin (Pdpn), Prox1, and Sox18 individually (Figure 3C). All tgKSHVΔRta tumors robustly transcribed CD34, Flt4/VEGFR-3, Lyve1, and Sox18. Pdgfra, Pdpn, and Prox1 mRNAs were detectable as well. This demonstrates that the murine KS tumors exhibit a LEC-specific transcription profile similar to human KS. The tumors that developed in tgKSHVΔRta were similar (Figure 3D, E). Based on principal component PC1 loadings, three pathways influenced subtypes: blood (Hba-a2), muscle (Ckm, Tnnc2, Acta1), and B cells (Jchain, Ccl19, Glycam1, Cd79a) (Figure 3G). PC2 loadings were dominated by mRNAs expressed in salivary gland tissue: Smr3a, Chia1, Cst5, Glycam1, Amy1, Bpifb1, Atp2a1 (Figure 3H). These data are consistent with the tumors originating in the head and neck region, near or invading salivary glands and cervical lymph nodes.

**Figure 3:**
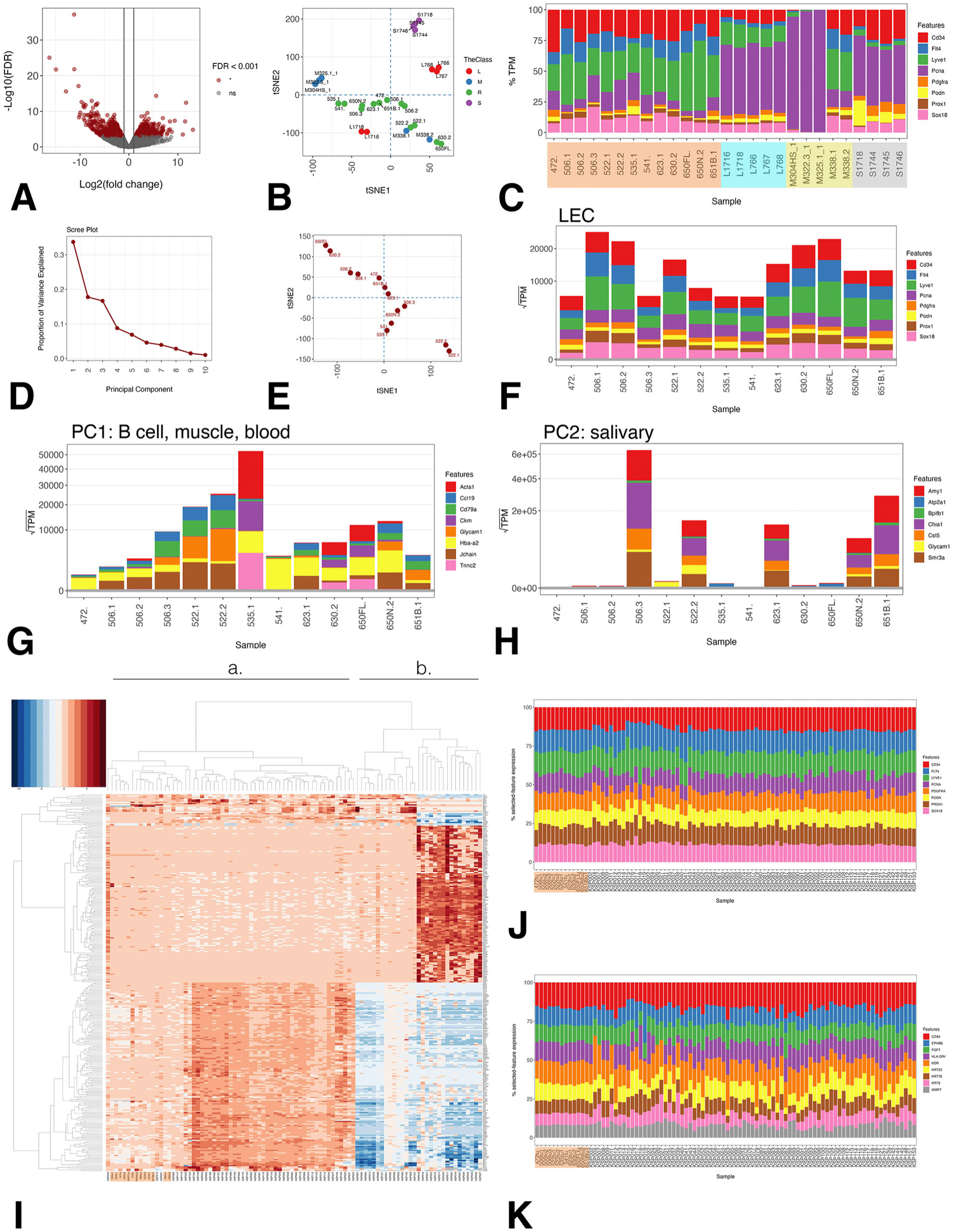
Host transcription. (A) Volcano plot of 13 tumors from tgKSHVΔRta vs 5 from tgKSHVΔmiR. FDR rate is on the vertical axis and log2 fold change on the horizontal axis. Differentially regulated genes (FDR-adjusted p≤0.001) in red. (B) tSNE projection of all samples based on differential transcription. Lymph nodes “L” in red, tgKSHVΔmiR tumors “M” in blue, tgKSHVΔRta tumors “R” in green, and salivary glands “S” in purple. (C) %TPM for EC lineage markers, PCNA, and PDGFRα. Salivary glands in gray, tgKSHVΔmiR in green, lymph nodes in cyan, and tgKSHVΔRta in orange. (D) Scree plot of proportion of variance explained of tgKSHVΔRta tumors and (E) tSNE. (F) TPM for EC markers. (G, H) TPM of genes contributing significantly to PC1 and PC2 (I) Heatmap of RNAseq for human KS and tgKSHVΔRta murine KS (two significant branches and four clusters), tgKSHVΔRta tumors in orange. (J) %TPM for EC lineage markers, PCNA, and PDGFRα and (K) other lineage markers across mouse and human tumors, tgKSHVΔRta in orange.

Next, we compared human and murine KS (Figure 3I). Two main clusters were evident, corresponding to the two types of KS previously reported ^8^. Cluster **a** comprised both human and mouse samples; cluster **b**, only human samples. Key KS markers (CD34, Flt4/VEGFR-3, Lyve1, PDGFR-α (Pdgfra), Podoplanin (Pdpn), Prox1, and Sox18) were present at similar levels in both. Among the top 300 most variable mRNAs (Figure 3J), key skin markers (CD34, EphrinB6, FGF7 or Keratinocyte Growth Factor (KGF), HLA-DA/ MHCII, KDR / VEGFR-2, Keratin 23, Keratin 78, Keratin9, and MMP7) were detectable in human and murine KS (Figure 3K). The variation among individual lesions in human skin KS biopsies was greater than between murine and human.

### Tumors from tgKSHVΔRta mice transcribe putative KSHV oncogenes

Viral transcription in tgKSHVΔRta tumors was robust (Figure 4A). As expected, no reads mapped to the Rta gene. The E2F-promoter-driven GFP and Hygromycin**^r^** genes were constitutively transcribed. Transcription was highest on either side of the viral miRNA intron (Supplemental Figure 3A, B), at the two 3’-coterminal operons immediately adjacent to the TR (K1/ORF4 and K15/ORF75), and the K12/Kaposin, LANA, vCYC, vFLIP latency locus and the viral interferon regulatory locus, including vIRF3 or LANA2 ^24^. The herpesvirus-defining 3’-coterminal transcription cascade was maintained, as evidenced by a gradual increase in reads mapping to known multi-transcript-terminating poly (A) sites (Figure 4A, arrows), as were known intron/exon junctions (Supplemental Figure 3C). An independent analysis of batch-corrected TPM counts per ORF confirmed the analysis of raw reads (Figure 4B). Transcription was consistent across multiple independent tumors (Figure 4C). The K14 and ORF74/vGPCR mRNA, which is more strictly Rta-dependent than any other gene in the viral genome, ^25^ had very low and inconsistent expression. Late mRNAs that relied on ORF24, the viral TATA-binding protein that recognizes an aberrant KSHV-specific TATT element ^26^, were not detected above background. One could consider the entire late region ORF16 to ORF49 (43,000 bp), devoid of regulatory elements that respond to transcriptional activators in the absence of Rta.

**Figure 4:**
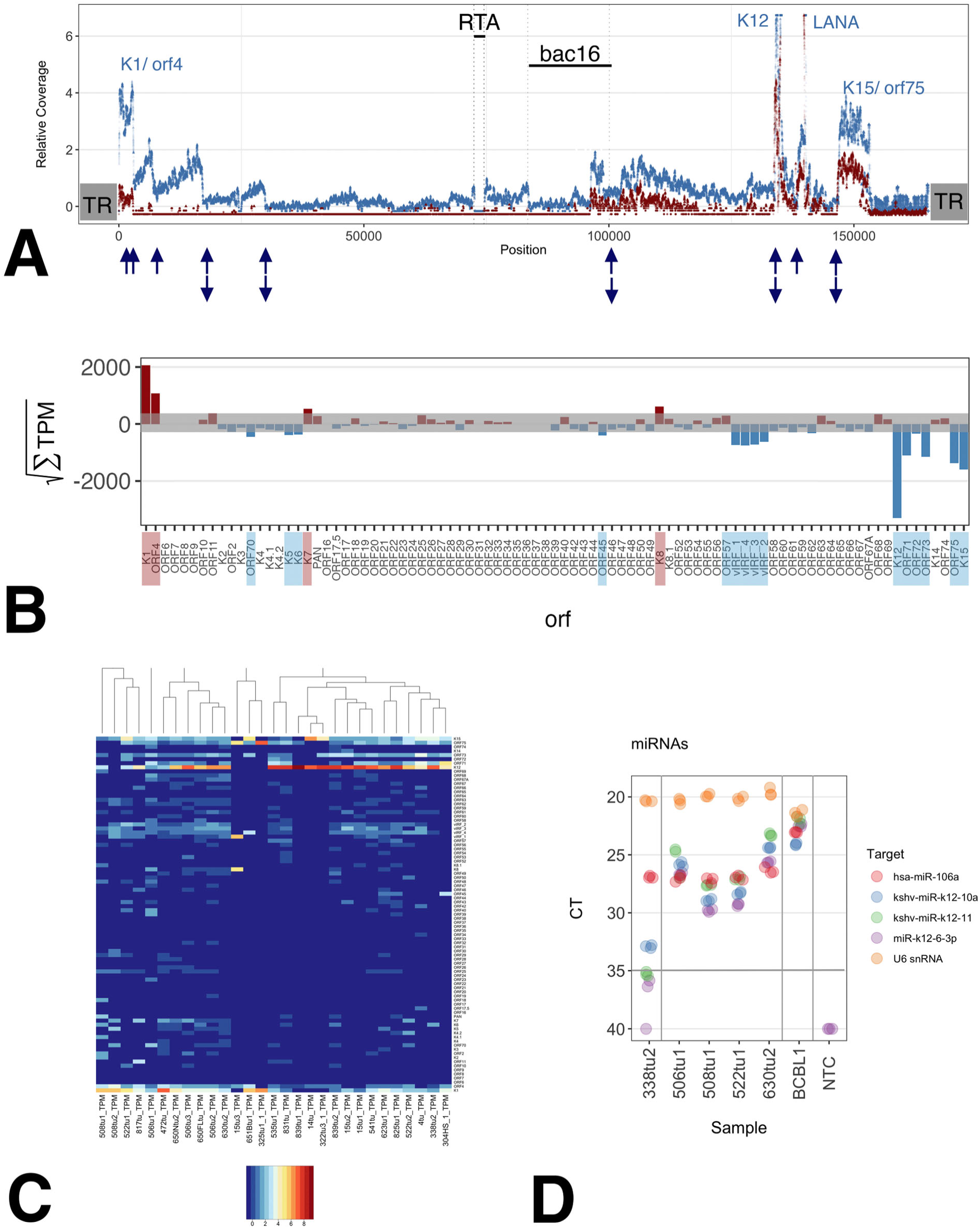
Viral transcription. (A) Relative coverage across the viral genome tgKSHVΔRta (blue) or tgKSHVΔmiR (red) tumors. Blue arrows indicate polyA sites. Terminal repeats (TR), bac16 (backbone), K1/ORF4 and K15/ORF75 (bi-cistronic transcripts). (B) Squared sum of TPM per ORF. Red indicates forward and blue reverse-oriented ORFs. (C) Heatmap. Samples are arranged by hierarchical clustering, ORFs by position. Red indicates higher and blue lower than the median overall count. (D) qPCR analysis for KSHV and host miRNAs. Raw CT values are on the vertical axis and samples are on the horizontal axis. Colors indicate different miRNA probes.

The viral miRNAs were expressed as determined by TaqMan real-time qPCR (Figure 4D). The abundant U6 snRNA marked the upper limit of the linear range. Normalized levels of viral miRNAs were lower in the tgKSHVΔRta tumors than in the KSHV-infected BCBL1 cell line; however, viral miRNA levels approached those of a conserved housekeeping miRNA, hsa-miR-106a, suggesting that the relative abundance of viral to host miRNA levels was equivalent and likely reflects genome copy number. This contrasted with the tgKSHVΔmiR tumors, where the deleted miRNAs represented by miR-K12-6-3p and miR-K12-11 were below the limit of detection, and miR-K12-10a was present at much reduced levels (miR-K12-10a, located within Kaposin was not deleted, but mutated, which affected qPCR primer binding efficiency). We conclude that the tgKSHVΔRta, as opposed to tgKSHVΔmiR mice, retained physiological levels of KSHV miRNAs, which are known to drive spindle cell differentiation.

### Cell lines from tgKSHVΔRTA tumors lose viral gene expression

In the presence of endogenous growth factors, primary tumor explants formed a monolayer of spindle-shaped cells (Supplementary Figure 5A). 3/12 (25%) of primary cultures could not be expanded (Supplementary Figure 5B, C). From 9/12 (75%) explant cultures, immortalized cell lines emerged, as is typical for rodent cells ^27^ (Figure 5G). The highest passage cell line, d5010tu630, lost contact inhibition, became growth factor-independent, and gained the ability to grow in soft agar (Figure 5D, E, F). This cell line, but not two others, formed tumors upon subcutaneous injection into NSG mice (Supplemental Figure 5H). LANA was expressed in the primary explant population (Supplemental Figure 5I), but expression was lost with passage (Supplemental Figure 5J). None of the cells remained GFP-positive or hygromycin-resistant, despite these markers being controlled by the E2F-houskeeping gene promoter. A similar loss of KSHV and KSHV gene expression is consistently seen in cells exogenously infected in culture ^28,29^. No human KS tumor-derived cell lines exist, suggesting that one or more viral genes are incompatible with current culture conditions for EC.

## Discussion

Transgenic mice are a powerful tool for tumor virology. The first transgenic mouse did not use a defined human oncogene but rather one half of the SV40 genome, comprising the native viral enhancer, promoter, and large T oncogene ^30^. The assumption, then and now, is that fundamental molecular mechanisms of tumorigenesis and gene regulation are evolutionarily conserved across species. This study confirms this general principle and extends it to KSHV.

The tgKSHVΔmiR and tgKSHVΔRta mice both developed LANA-positive vascular sarcomas that recapitulated human KS. The lesions appear disordered, invasive, and were often multifocal. KS is defined by its viral etiology and by the expression of LEC lineage markers, particularly VEGFR-3, LYVE-1, and Podoplanin ^7,9^. LECs uniquely enable KSHV gene expression to the point of tumor induction. KSHV has evolved to respond to the developmental cues in LEC. Prox18 is an obvious candidate to regulate both KSHV and LEC transcription ^31^. The tgKSHVΔRta mice constitute a robust and so far only *in vivo* model for investigating this interconnection. Targeting this synthetic dependence may yield tissue-, infected cell-, and tumor-selective therapies.

Tumors in tgKSHVΔRta mice were larger, more invasive, and more multifocal than those induced by ectopic overexpression of any single viral gene ^32^. This suggests that multiple KSHV genes cooperate to cause the signature pathology of KS. TgKSHVΔRta mice had a higher OS than tgKSHVΔmiR mice due to a lower incidence of angiectasis. Murine KS developed adjacent to the salivary gland and cervical lymph nodes, as is common in pediatric KS ^3^. The tumor phenotype was independent of mouse strain background.

The KSHV TR regions exhibited enhancer properties, as the TR-proximal promoters K1/ORF4 and K15/ORF75 were the most active. This *in vivo* finding extends a recent observation from cell culture ^33^. All known KSHV latent promoters, including those in the vIRF1-4 region, were active. Their activity is evolutionarily conserved between mouse and humans and independent of Rta. Some “classic” lytic genes, which are controlled by Rta-response elements, were active in the absence of Rta, indicative of multimodal regulation. This includes ORF75, which, until recently, was considered a lytic gene, but has since been detected in KS biopsies independent of lytic gene expression ^34,35^, and ORF K1, which is expressed during latency but highly induced upon virus reactivation ^36^. Notably, K1 can extend the lifespan of primary human ECs^37^. By contrast, transcripts controlled by an unconventional TATA box, which is PIC/ORF24-dependent ^26^, including the viral DNA polymerase, were not detectable. This is consistent with the notion that lytic viral DNA replication and viral structural proteins are not required for tumorigenesis, provided the viral genome persists by other means, such as being tethered by LANA to or integrated into the host chromosome. Many KS patients, even with untreated AIDS-KS, have no detectable virus in blood ^2,8^. For other herpesviruses, too, lytic DNA replication is not required for lifelong persistence ^38^. This experiment separates KSHV’s transforming activity from its lytic DNA replication and virion-producing functions.

A fraction of the tgKSHVΔRta mice developed lymphoid hyperplasia. This pathology mimics the natural history of KSHV. In addition to KS, KSHV causes B-cell hyperplasia, multicentric Castleman’s disease (MCD), and PEL (reviewed in ^39^). Many AIDS-KS patients present with concurrent lymphoma or MCD ^40^. Our prior LANA/vCyc/vFLIP/miRNA KSHV transgenic mice were hyperresponsive to antigen and developed frank DLBCL when crossed in Myc-mutated mice ^41^. Other individual KSHV proteins, vIL6 and vCYC, also induce lymphoma in transgenic mice (reviewed in ^42^). With the tgKSHVΔRta mice, we now have an immunocompetent model to study the co-occurrence of lymphoid and solid-organ tumors associated with KSHV and their dependence on immune-cell signaling.

### Limitations of the study

This model, by design, is not dependent on viral replication since the tgKSHVΔRta transgene lacks Rta and does not express the viral polymerase or capsid genes. It is an advantage as the mice can be co-housed and interbred without enhanced biosafety requirements for human infectious agents. It is a limitation, as we know KS to be a disseminated disease initiated by virus reactivation after decades of molecular latency. This reactivation event requires Rta. Reports on defective KSHV genomes ^43^ suggest that some KS skin lesions may result from metastatic cell spread and not depend on viral reactivation. By contrast, studies on KS tumor heterogeneity suggest that most skin lesions are polyclonal and result from independent infection events ^12^. In humans, LANA is present in 100% of KSHV-infected spindle cells and the viral episome is necessary for the cells to survive ^19,44^. Future studies are needed to test whether LANA becomes dispensable if KSHV is integrated into the host genome.

As with all cancers ^8^, we can only measure the genes that are transcribed in established tumors, not those that are only transiently required. The vGPCR (ORF74) was inconsistently transcribed at low levels in tgKSHVΔRta mice, consistent with mRNA and protein not being detectable in KS lesions or latent PEL; however, when expressed in isolation, vGPCR functions as a potent transforming gene in culture and *in vivo*^32^. There may be a developmental stage, perhaps at the stem cell or progenitor state, at which vGPCR is required for KSHV pathogenesis; otherwise, it would not have remained functional under purifying selection. Further experiments are needed to identify these minor cell populations.

KS tumors contain a complex mixture of cell types. They do not exhibit signature host mutations ^12^. By scRNA-seq, KSHV mRNAs were detected primarily in LEC in human KS lesions ^9^. By spatial proteomics and conventional histochemistry, LANA colocalizes with VEGFR-3 and LYVE-1 ^7^. These experiments, however, cannot prove where tumor-resident LECs originate or whether they are trans-differentiated mesenchymal cells that express LEC markers. ScRNAseq cannot prove cells that do not express KSHV mRNAs in a KS biopsy, never did, or do so at a level below the limit of detection, as more abundant mRNAs are always more likely to be detected over rare ones. tgKSHVΔRta opens the possibility of overcoming the limitations of descriptive analyses through targeted genetic manipulations of the host.

## Acknowledgments

We gratefully acknowledge the late Sang-Hoon Sin for initiating this work and for their invaluable contributions to the project.

This project was funded by Public Health Service grant 5R01CA250080-05. K.W.S. was supported by T32 2T32AI007419-31.

We thank the UNC Animal Models Core, Preclinical Research Unit, and Pathology Services Core, whose work is supported in part by NCI Center Core Support Grant P30CA016086.

## Methods

### Bacterial Artificial Chromosome Cloning

Using a Red/ET recombineering cloning strategy utilizing the pRedET plasmid and primers H1-RN-N and H2-RN-C, the ORF50 CDS (2,058 bp) of the full-length KSHV genome on a bacterial artificial chromosome (BAC16)^18^ was replaced with a RpsL-Neomycin cassette. BAC16 ORF50/Rta deletion (BAC16ΔRta) clones were then selected via kanamycin resistance, and purified BAC16ΔRta plasmid was confirmed using Kpn1 restriction digest and pulse-field gel electrophoresis. To confirm the deletion of ORF50, the BAC16ΔRta plasmid was subjected to Ion Torrent sequencing on the Ion GeneStudio S5 System (ThermoFisher), with gap regions filled via PCR amplification-based Sanger sequencing through Genewiz (Azenta Life Sciences). PCR amplification of gap regions was performed using primer sets Gap 1-6. The BAC16ΔRta consensus sequence is publicly available on GenBank (MN688631.1).

### BAC Transfection and Cell Line Creation

Stable BAC16-carrying cell lines were established as previously described^45^. HEK293T cells were transfected with BAC16 or BAC16ΔRta DNA, selected with hygromycin treatment, then grown in co-culture with iSLK cells carrying a doxycycline-inducible RTA construct (iSLK-RTA). After selection of all constructs with G418, Puromycin, and Hygromycin, doxycycline and sodium butyrate were added to the culture to induce lytic reactivation and replication. The resulting recombinant KSHV virus was then used to infect iSLK-RTA cells, which were again selected with G418, Puromycin, and Hygromycin to create the stable cell lines ΔRTA-BAC16-iSLK and WT-BAC16-iSLK. These stable BAC16-iSLK-RTA cell lines were then induced using doxycycline and sodium butyrate to produce recombinant KSHV virus, which was in turn used to infect HEK293 cells. Infected HEK293 cells were selected with hygromycin to create the BAC16-293 stable cell lines ΔRTA-BAC16-293 and WT RTA-BAC16-293. For western blotting of stable cell lines in lytic reactivation conditions, 293 stable cell lysates were collected 48 and 72 hours post-induction with sodium butyrate and TPA (12-O-tetradecanoylphorbol-13-acetate), while the BAC16-iSLK-RTA cell line lysates were collected immediately before and 48 hours after induction with sodium butyrate and doxycycline.

### Transgenic Mouse Line Creation and Maintenance

BAC16ΔRta was purified using a QIAfilter Plasmid Midi Kit (Qiagen, 12243) for pronuclear injection of FVB/NJ mouse oocytes. Pronuclear injections and subsequent implantations to generate transgenic mice were performed by the University of North Carolina at Chapel Hill’s Animal Models Core Facility. Transgenic founder mice were bred with FVB/NJ mice to produce an F1 generation of transgenic mice, then transgene-positive brother-sister mating pairs were used to maintain the tgKSHVΔRTA mouse line, which was submitted to MMRRC for cryopreservation at generation F11. Mouse genotyping was performed using DNA extracted from mouse tail clippings using the Maxwell RSC Genomic DNA Kit (Promega, AS1880) on the Maxwell RSC instrument (Promega) and PCR primers targeting LANA, vFLIP, RTA, KSHV pre-miRNA, and murine ApoB. Mouse lines were housed in hot-wash cages in a facility accredited by AAALAC International. All experiments were approved by UNC’s Institutional Animal Care and Use Committee (IACUC) # 24-152 and 25-251.

### Histology and Immunohistostaining

Tissues were fixed in 10% neutral buffered formalin for 24-72 hours, then exchanged to 70% ethanol and kept at 4°C until embedding. All tissues underwent paraffin embedding, sectioning to 5 µm and staining with hematoxylin & eosin (H&E) via the Pathology Services Core of the UNC Lineberger Comprehensive Cancer Center. Tissue slides were deparaffinized and rehydrated using a gradient of Histochoice Clearing Agent (Millipore-Sigma, H2779-1L), ethanol, and water on the Leica ST4020 linear stainer (Leica). Slides underwent antigen retrieval (AR) in the PT Link instrument (Agilent) using AR buffer (10 mM citrate buffer, 0.05% Tween-20, pH 6.0) for 25 minutes at 95°C. Slides were washed for 5 minutes in 1X Dako wash buffer (Agilent, S300685-2C) then used for either IHC or IFA.

IHC staining was performed by treating tissues with 3% hydrogen peroxide for 5 minutes, followed by serum blocking (0.1% gelatin, 0.5% BSA, 5% animal serum, 0.1% Triton X-100, 0.05% Tween-20, in PBS) for 20 minutes, followed by avidin and biotin blocking solutions (Vector Labs, SP-2001) for 15 minutes each, with two 3-minute PBS washes between each blocking step. Serum for blocking was from the same species as the secondary antibody. After a PBS wash, each slide was incubated overnight at 4°C with primary antibody diluted in diluent (0.1% gelatin, 0.5% BSA, 0.1% Triton X-100, in PBS) using one of the following: anti-LANA (1:75) (rabbit) clone ZR106 (Zetacorp, Z2531RS), anti-VEGFR-3 (1:100) (rat) clone AFL4 (Invitrogen, 14-5988-82), anti-Ki-67 (1:100) (rabbit) clone SP6 (Epredia, RM-9106-SO), anti-LYVE-1 (1:100) (rat) clone ALY7 (Novus Biologicals, NBP1-43645), or no primary antibody control. Human KS tissues for IHC were stained with clinical LANA antibody anti-LANA (1:500) (mouse) clone 13B10 (Leica, NCL-HHV8-LNA). After 3 washes in PBS, tissues were incubated for 10 minutes in secondary antibody solution (5% animal serum, HRP-conjugated secondary antibody [1:100], in PBS), followed by 3 more washes in PBS. Tissues were then incubated for 5 minutes in avidin/biotin complex solution (4% Reagent A, 4% Reagent B in PBS, made 30 minutes before use). HRP-conjugated secondary antibodies, blocking serum, and the avidin/biotin complex solution reagents sourced from VECTASTAIN Elite ABC (species-specific) IgG kit(s) (Vector Labs, PK-6101, or PK-6104). After 3 washes with PBS, tissues were incubated with ImmPACT NovaRED (HRP) substrate solution (Vector Labs, SK-4805) for 5 minutes. Slides were then quenched in ultrapure water, stained briefly with Mayer’s Hematoxylin (Electron Microscopy Sciences, 26043-05) followed by 1:100 ammonium hydroxide. Slides were immersed for 5 minutes in running tap water before dehydration with a gradient of ethanol to Histochoice on the linear stainer. Slides were cleared in xylene before mounting with Shandon-mount (Epredia, 1900331) and coverslip. Stained slides were viewed on a Leica DM4000B microscope (Leica Microsystems) with objective lenses of 10x, 20x, or 40x magnification, and images were captured on an Axiocam105 color camera via Zen imaging software (Zeiss).

IFA staining was performed by first incubating AR-treated tissue slides in permeabilization buffer (0.5% Triton X-100 in TBS) for 10 minutes and washing twice in TBS-T (0.1% Tween-20 in TBS) for 5 minutes each. Slides were then incubated in 2.5% normal horse serum (NHS) for 20 minutes; pre-diluted serum was obtained from VectaFluor Excel Amplified Kit - Anti-Rabbit IgG DyLight 594 (Vector Labs, DK-1594). Each slide was incubated for 2 hours with either anti-LANA primary antibody (1:75) clone ZR106 (Zetacorp, Z2531RS) or no-primary diluent (5% NHS in TBS-T), followed by two 5-minute TBS-T washes. VectaFluor kit components were used for a 15-minute incubation with goat anti-rabbit secondary, followed by 30 minutes with horse anti-goat DyLight 594 tertiary; two 5-minute TBS-T washes were done between and after the VectaFluor reagents. Tissues were stained with DAPI (10 µg/mL in TBS) for 1 minute, washed twice in TBS, and mounted with glass coverslips using ProLong Glass Antifade Mountant (Invitrogen, P36982). After curing for 24 hours, slides were imaged on the Zeiss Axio Observer 7 fitted with a 63x oil objective, Apotome 3 structured illumination microscopy (SIM) filter, and Axiocam 807 mono microscope camera. Images were captured in Zen 3.11 software (Zeiss) utilizing z-stacks deconvoluted with Apotome Plus (Zeiss) and converted from SIM optical sections to a maximum intensity projection.

### Long Read PacBio DNA Sequencing

For evaluation of the transgene insertion site, spleens from transgene-positive homozygous mice were extracted whole and immediately frozen at −20°C. Frozen spleens were diced into small pieces, then 10 mg of tissue was digested with proteinase K and treated with RNase, then DNA was extracted using the DNeasy Blood and Tissue Kit (Qiagen, 69504). Before sequencing, DNA samples were analyzed for concentration and quality utilizing the Nanodrop 1000 (ThermoScientific), Qubit Flex Fluorometer (Invitrogen-ThermoFisher Scientific), and 4200 Tape Station (Agilent Technologies). Genomic DNA was sheared using Covaris g-TUBEs (Covaris, 520079), following the PacBio technical note “Covaris g-Tube DNA Shearing for SMRTbell Prep Kit 3.0”. DNA libraries were created using the SMRTbell Prep Kit 3.0 (Pacific Biosciences, 102-141-700), following manufacturer instructions with the modification of increasing the 37°C A-tailing thermocycler step from 30 minutes to 45 minutes. The Sequel II Binding Kit 3.2 (Pacific Biosciences, 102-194-100) and Sequel II Sequencing Kit 2.0 (Pacific Biosciences, 101-820-200) were used as per manufacturer instructions to prep, load, and sequence the DNA libraries on the Sequel IIe instrument (Pacific Biosciences).

For insertion site determination, the “Map Long Reads to Reference” tool in CLC Genomics Workbench (Qiagen) was used: PacBio HiFi reads were first mapped to the KSHV BAC16 reference genome (GenBank MK208323.1), then all reads that mapped to KSHV BAC16 were then mapped onto the mouse reference genome (Mus musculus C57BL/6 GRCm39 (mm39, PRJNA20689)) to find reads that covered both the mouse genome and the KSHV transgene.

### RNA Extraction from Tissue

Extracted tissues were placed into RNAlater (Invitrogen, AM7022) according to manufacturer instructions, then frozen at −80°C for long-term storage. RNA was extracted using the Maxwell RSC miRNA Tissue Kit (Promega, AS1460) on the Maxwell RSC instrument (Promega). Immediately before extraction, 10-15 mg of thawed and dried tissue from each sample was homogenized in tubes with 2.38 mm beads (Qiagen, 13117-50) on the TissueLyser II instrument (Qiagen). RNA quality and quantity were measured using the Nanodrop One spectrophotometer, Qubit Flex fluorometer, and Tapestation 4200 instruments.

### Bulk RNA-sequencing

For bulk RNA-seq, sample poly-A (mRNA) isolation, RT-PCR, library prep, and subsequent sequencing on the Illumina NovaSeq X Plus (25B lane, 2×150bp) was performed by Genewiz (Azenta Life Sciences), as was the sequencing data demultiplexing, QC, and adapter trimming. Using the RNA-Seq Analysis tool in CLC Genomics Workbench (Qiagen), trimmed reads were then mapped to the mouse reference genome (Mus musculus C57BL/6 GRCm39 (mm39)), then all unmapped reads from that mapping were subsequently mapped to the KSHV BAC16 reference genome (GenBank MK208323.1). Transcription levels across the TRs could not be evaluated as the short reads did not map uniquely. Reads that mapped to the TR, OriLyt, and LANA repeat regions were discounted, as their mapping was not unique and contaminated with reads derived from host repeats.

Coverage analysis of KSHV. Reads were aligned to MK208323 the sequence of the JSC1-derived BAC16 bacmid as determined at City of Hope. To calculate the sum total across all tumors, the raw mapped reads at each position of the genome were tabulated, and variance was stabilized by Anscombe transformation, which is less radical than the logarithmic transformation. A floor of 3 reads and a ceiling of 100 reads per position was introduced.

Isolation Forest. Outlier detection was performed using the Isolation Forest algorithm implemented in the H2O package. The algorithm identifies anomalies by constructing an ensemble of isolation trees that recursively partition the data using random feature and split-value selection. Anomalous samples require fewer splits to isolate and thus have shorter average path lengths across trees. The model was trained on 15 samples from tgKSHVΔRta tumors, with 13,691 gene features, using 100 trees and default parameters. Samples in the bottom 10th percentile of anomaly scores were flagged as potential outliers for further investigation.

Comparison of human and mouse transcription. As human and mouse RNA-seq datasets were generated using different protocols and sequencing depths, raw counts in each dataset were normalized independently using DESeq2’s median-ratio size-factor method, which is more robust than TPM. Next a variance-stabilizing transformation (VST) was applied. The 2,000 most variable Human–mouse orthologs were selected for downstream analyses. Hierarchical clustering was performed using Ward.D2 linkage on Euclidean distances. There are some limitations to this analysis. Human KS skin punch biopsies are highly variable and guided by clinical considerations to limit bleeding and discomfort. They encompass varying amounts of tumor and normal skin, whereas in the mouse tumors could be cleanly dissected out. We therefore removed the human KS cases with the highest keratin mRNA levels, indicative of differentiated skin epithelial cells.

### KSHV miRNA qPCR Assay

10 ng of RNA from 17 tumor and 8 normal tissue samples was reverse-transcribed using the TaqMan™ MicroRNA Reverse Transcription Kit (ThermoFisher Scientific, 4366596), utilizing the target-specific stem loop primer in each of the following TaqMan MicroRNA Assays; kshv-miR-k12-6-3p (008459), kshv-miR-k12-10a (008504), kshv-miR-k12-11 (008562), hsa-miR-106a (002169) (ThermoFisher Scientific). The resulting cDNA, primers and probes from the associated kits, and TaqMan™ Universal PCR Master Mix, no AmpErase™ UNG (ThermoFisher Scientific, 4324018), were used in a qPCR reaction to detect each target. QPCR was run on a QuantStudio 7 Pro (ThermoFisher Scientific).

### Establishment of Mouse Tissue-derived Cell Lines

Tumor tissue or extracted lymph nodes were cut into pieces then placed into a gentleMACS C tube (Miltenyi Biotec, 130-093-237) in DMEM (Gibco, 11995-065) containing 0.25% Collagenase Type 2 (Worthington Biochem, LS004202). Each C tube was subjected to mechanical griding using program m_impTumor_02 on the gentleMACS Dissociator instrument, then incubated for 40 minutes at 37°C on a rotator, followed by another mechanical disruption using program m_impTumor_03 on the Dissociator instrument. The resulting slurry was passed through a 100 µm cell filter mesh (Fisher Scientific, 22363549), with any remaining solids ground through the filter with a 1 mL syringe plunger, then went through 2-3 rounds of RBC lysis (Sigma, R7757). The whole tumor cell pellet was resuspended in 1 mL Endothelial Cell Growth Media 2 (PromoCell, C-22111), with 10% fetal bovine serum (FBS) (Corning, 35-015-CV) and all growth factors added except heparin, then plated in a 35 mm cell culture dish (Greiner Bio-One, 627102). After 72 hours the attached cells were trypsinized and transferred to a T25 cell culture flask (Genesee Scientific, 25-207) where they were grown in ECGM2 and split 1:1 or 1:2 every 3-5 days until transformation.

### Soft Agar Colony Forming Assay

Autoclave sterilized 6% agar (BD, 214220) in ultrapure water was melted and cooled to 42°C in a bead bath, then mixed with 42°C ECGM2 media 1:10 to create 0.6% agar ECGM2 media. 0.5 mL of 0.6% agar ECGM2 was added to each well of a 24 well plate (Falcon, 353047), creating a base layer. After the base layer had solidified and cooled, 6% agar was heated and cooled to 42°C, then mixed with 42°C ECGM2 media 1:15 to create 0.4% agar ECGM2. The 0.4% agar ECGM2 was immediately added to a freshly spun down cell pellet, mixed quickly, then 0.5 mL was dispensed on top of the 0.6% agar ECGM2 base layer in each well of the 24 well plate. Each cell line was plated at three concentrations: 2000, 1000, and 500 cells per well, in duplicate. After 12-14 days, the wells were imaged on a Leica MZ6 surgical scope (Leica Microsystems) and individual colonies were counted manually looking through the 4x objective on a Leica DMIL microscope (Leica Microsystems, 090-135.001) and imaged using an Axiocam ERc5s (Zeiss).

### Tumor Allograft in NSG mice

NSG mice (NOD.Cg-*Prkdc^scid^ Il2rg^tm1Wjl^*/SzJ), female, were injected subcutaneously in the right rear flank with 1×10^6^ cells in a 1:1 mixture of PBS and Matrigel, 200 µL total per mouse. Injected cell lines included 630 (passage 95), 685 (passage 79), and 690 (passage 56), n = 5 mice per cell line. Mice were monitored regularly for health and tumor growth and were collected when the first tumor exceeded 1 cm in size, 43 days post-injection.

## Supplemental Figures

**Supplemental Table 1**

Comparison of KSHV founder animals that were generated in this study using BAC16 KSHVΔRta and previously using wild-type BAC16 KSHV and BAC16 KSHVΔmiR.

**Supplemental Table 2**

Pathologist evaluation of 31 tumors from n = 17 tgKSHVΔRta mice.

**Supplemental Figure 1.**

Histological comparison between human KS and tgKSHVΔRta murine KS. (A) H&E stain and (B, C) LANA stain of tgKSHVΔRta tumor with features resembling lymphangioma-like KS at two different magnifications (mouse #541). (D) H&E stain and (E, F) LANA stain of human KS presenting features of lymphangioma-like KS at two different magnifications. (G) H&E stain and (H, I) LANA stain of tgKSHVΔRta tumor within a cervical lymph node at two different magnifications (mouse #472). (J) H&E stain and (K, L) LANA stain of human lymphadenopathic KS at two different magnifications. (M) H&E stain of tgKSHVΔRta tumor infiltrating muscle (mouse #541). (N,O) H&E stain of tgKSHVΔRta murine KS in skin at two different magnifications (mouse #730). (P) H&E stain of human KS spindle cells infiltrating muscle (Q, R) H&E stain of human KS in skin at two different magnifications. For all panels, grey bars = 100 µm, black bars = 50 µm, and white bars = 20 µm.

**Supplemental Figure 2.**

Lymphocytes in tgKSHVΔRTA tumors. (A) H&E staining, (B) VEGFR-3, (C) Ki-67, and (D) no primary antibody control IHC staining of a tgKSHVΔRta tumor (mouse #535) within a lymph node showing lymphocyte proliferation at the tumor/lymph node interface (E) H&E staining, (F) VEGFR-3, (G) Ki-67, and (H) no antibody control IHC staining of a tgKSHVΔRta tumor (mouse #506) within a lymph node showing expanded germinal centers with proliferating lymphocytes. (I, M) H&E staining showing expanded germinal centers in lymphoid tissue within a murine KS tumor at two different magnifications (mouse #623). (J, N) H&E staining showing expanded germinal centers within a lymph node associated with a murine KS tumor at two different magnifications (mouse #651). (K, O) expanded germinal centers within a lymph node distal to the tumor site at two different magnifications (mouse #651). (L, P) extravascular erythrocytes within a lymph node distal to the tumor site at two different magnifications (mouse #650). For all panels, grey bars = 100 µm, black bars = 50 µm, and white bars = 20 µm.

**Supplemental Figure 3.**

Canonical viral transcription from the tgKSHVΔRta transgene. (A) RNAseq reads from 13 tgKSHVΔRta tumors mapped to KSHV BAC16 show highest read density is in unpaired reads (lighter colors) on either side of the viral miRNA locus intron. (B) When the intron is removed from the reference sequence, the aligned reads become paired (darker colors), showing the highest density of reads match a transcript from which the viral miRNA intron is spliced out. (C) RNAseq reads mapped to the BAC16 reference sequence show canonical splicing of K15, a viral gene with well-defined splice sites.

**Supplemental Figure 4.**

Establishment of tumor-derived murine KS cell lines. Tumor cells from a tgKSHVΔRta murine KS tumor (#630) (A) after zero passages (48 hrs after initial plating), (B) after 1 passage, (C) after 18 passages, and (D) after 48 passages. (E) Colonies formed in 24 well plate by 630 cell line during soft agar colony forming assay and (F) close up of the individual colonies. (G) Cell lines derived from tgKSHVΔRta mice and their current maximum passage number, with red bars representing tumor-derived lines from transgene positive mice and blue bars representing lines established from the cervical lymph nodes of transgene-negative FVB littermates. (H) Subcutaneous flank injection of NSG mice with cell line 630 (passage 95), forming tumors in 5/5 mice after 43 days. (I) LANA IFA demonstrating LANA expression in low-passage tumor-derived tgKSHVΔRta cell lines (white bars = 20 µm). (J) LANA IFA demonstrating lack of LANA expression in high-passage tumor-derived cell lines (white bars = 100 µm).

