## Supplemental Figures and Tables for "Kaposi Sarcoma herpesvirus reactivation and replication are dispensable for cell-lineage specific tumorigenesis in whole genome transgenic mice"

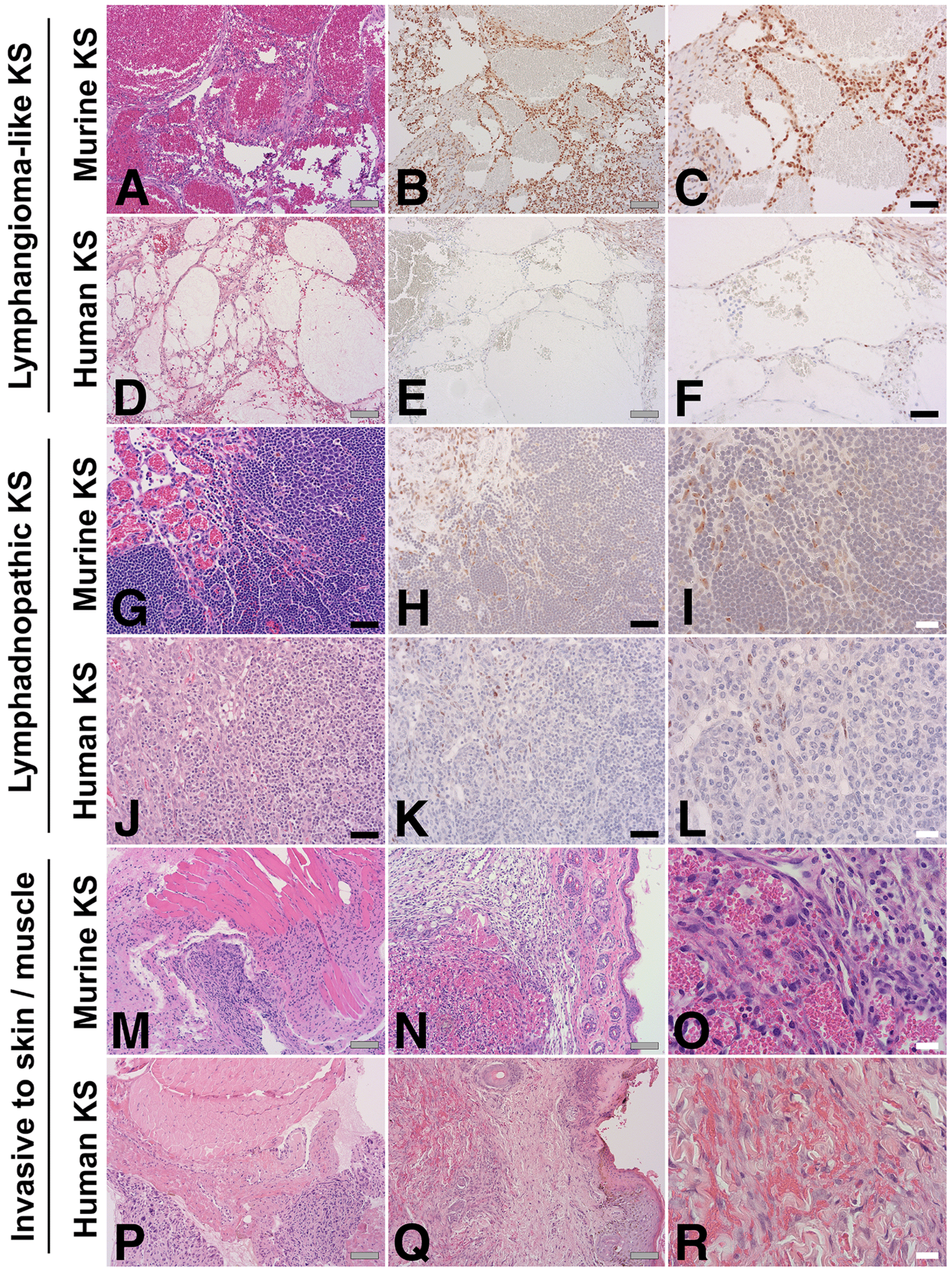
Lymphangioma-like KS
Murine KS
Human KS
Lymphadnopathic KS
Murine KS
Human KS
Invasive to skin / muscle
Murine KS
Human KS
A
B
C
D
E
F
G
H
I
J
K
L
M
N
O
P
Q
R

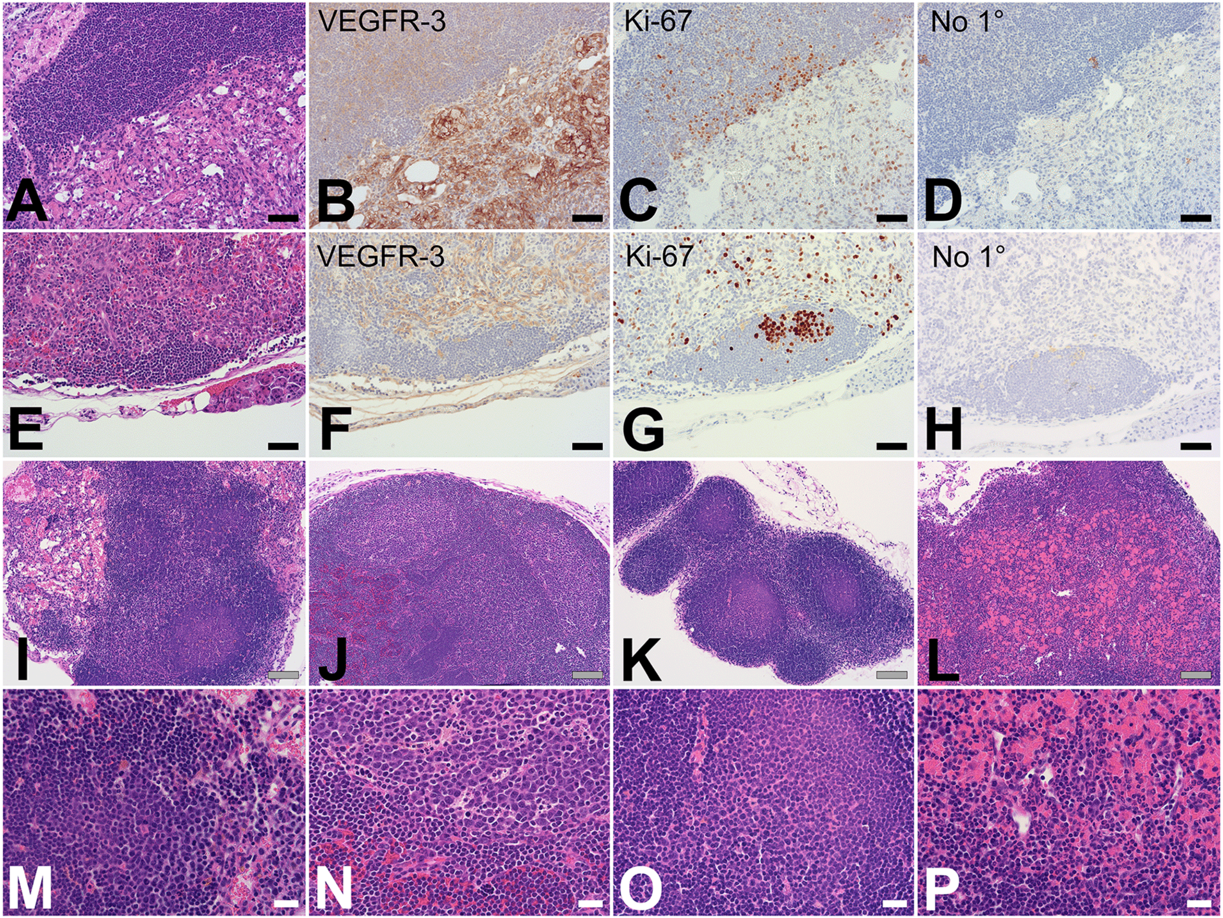
VEGFR-3
Ki-67
No 1°
VEGFR-3
Ki-67
No 1°
A
B
C
D
E
F
G
H
I
J
K
L
M
N
O
P

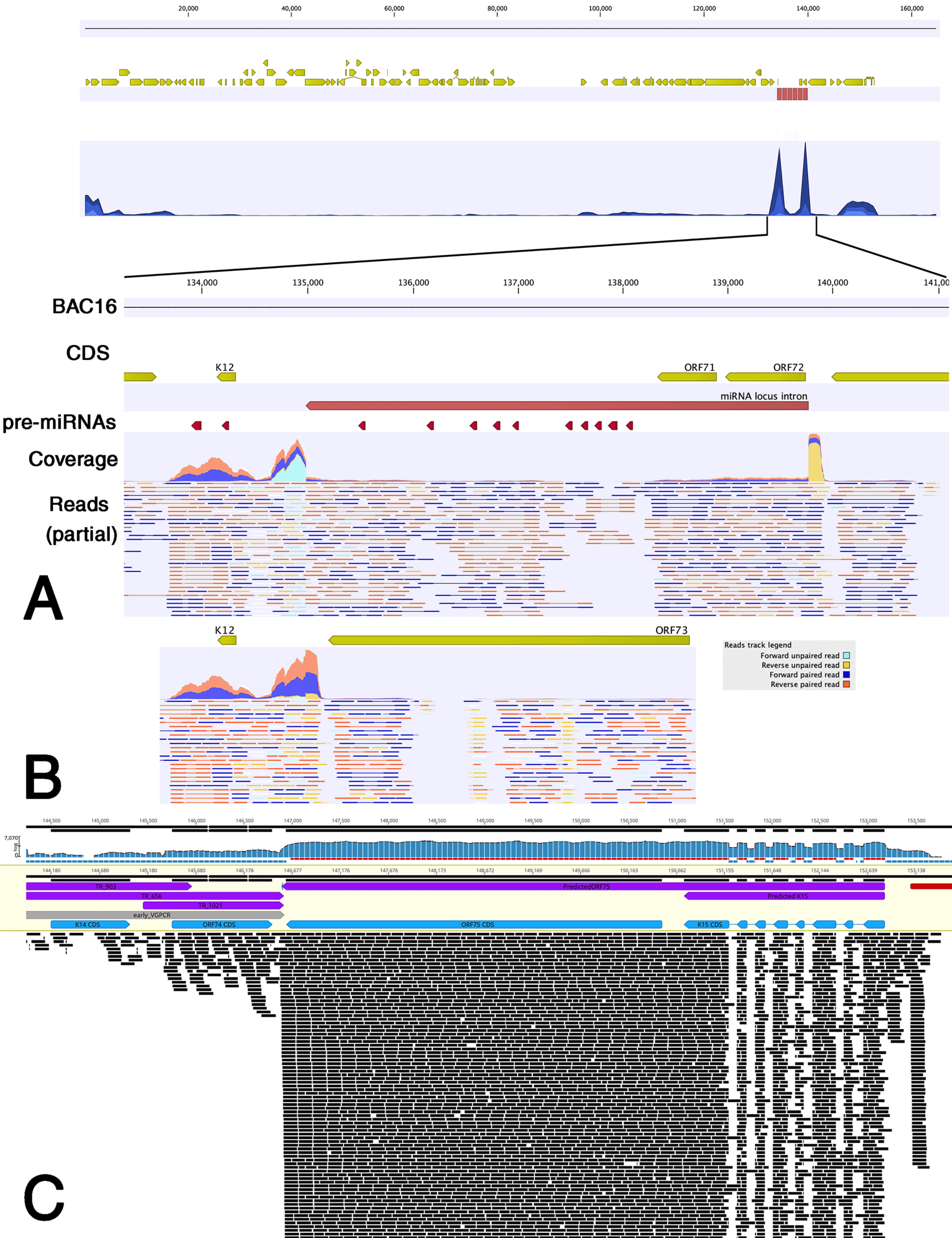

20,000
40,000
60,000
80,000
100,000
120,000
140,000
160,000
134,000
135,000
136,000
137,000
138,000
139,000
140,000
141,00
BAC16
CDS
K12
ORF71
ORF72
miRNA locus intron
pre-miRNAs
Coverage
Reads
(partial)
A
K12
ORF73
Reads track legend
Forward unpaired read
Reverse unpaired read
Forward paired read
Reverse paired read
B
TR_903
TR_656
TR_1021
early_VGPCR
K14 CDS
ORF74 CDS
PredictedORF75
ORF75 CDS
Predicted K15
K15 CDS
C

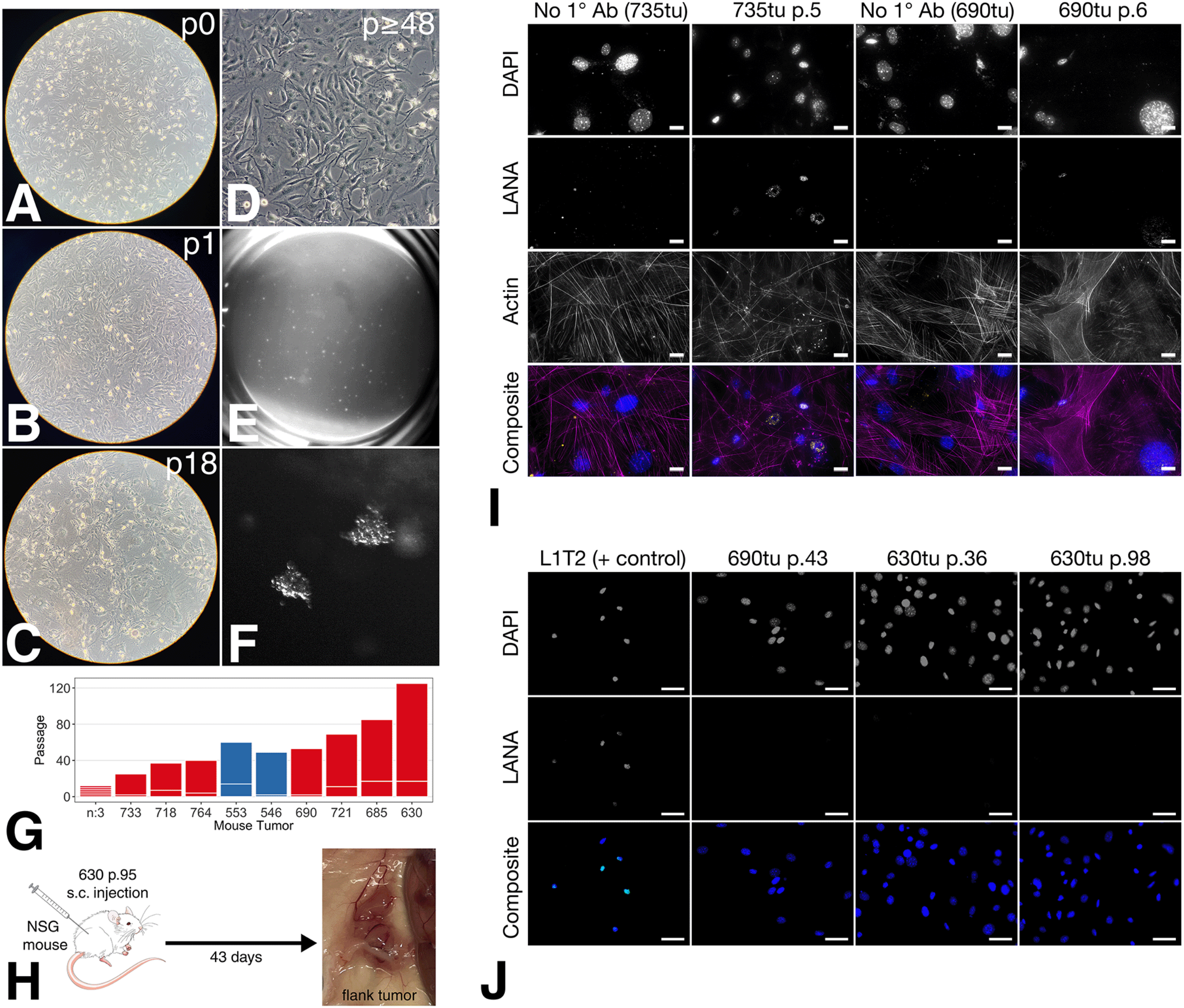

p0
A
p≥48
D
p1
B
E
p18
C
F
120
80
40
0
Passage
n:3
733
718
764
553
546
690
721
685
630
Mouse Tumor
G
630 p.95
s.c. injection
NSG
mouse
43 days
flank tumor
H
No 1° Ab (735tu)
735tu p.5
No 1° Ab (690tu)
690tu p.6
DAPI
LANA
Actin
Composite
I
L1T2 (+ control)
690tu p.43
630tu p.36
630tu p.98
DAPI
LANA
Composite
J

**Table 1:** Summary of KSHV-transgene founders

| Transgene (1) | Source | Injection (3) | F0 tag# | sex | F1 | strain | F0 age | F0 tumor | F1 tumor |
|---|---|---|---|---|---|---|---|---|---|
| BAC16 | pub (2) | a | KS.193 | M | no | FVB/N | 302 | no | . |
| | pub (2) | a | KS.194 | M | no | FVB/N | 239 | **yes** | . |
| | pub (2) | b | KS.1 | F | no | C57BL/6 | 26 | . | . |
| | pub (2) | b | KS.5 | F | no | C57BL/6 | 53 | **yes** | . |
| | pub (2) | b | KS.10 | M | no | C57BL/6 | 28 | **yes** | . |
| | pub (2) | c | KS.4 | M | no | FVB/N | 23 | **yes** | . |
| | pub (2) | c | KS.6 | F | no | FVB/N | 147 | no | . |
| | pub (2) | c | KS.8 | M | no | FVB/N | 9 | . | . |
| | pub (2) | c | KS.14 | M | no | FVB/N | 21 | . | . |
| BAC16ΔMIR | pub (2) | a | D.195 | F | no | FVB/N | 311 | no | . |
| | pub (2) | a | D.196 | F | no | FVB/N | 53 | **yes** | . |
| | pub (2) | a | D.197 | M | **Yes** | FVB/N | 100 | no | **yes** |
| | pub (2) | b | D.1 | F | no | C57BL/6 | 301 | no | . |
| | pub (2) | b | D.3 | F | no | C57BL/6 | 121 | no | . |
| | pub (2) | b | D.15 | M | no | C57BL/6 | 60 | no | . |
| | pub (2) | b | D.25 | M | no | C57BL/6 | 59 | no | . |
| | pub (2) | b | D.26 | M | no | C57BL/6 | 271 | no | . |
| | pub (2) | c | D.5 | F | no | FVB/N | 26 | . | . |
| | pub (2) | c | D.6 | F | no | FVB/N | 26 | no | . |
| BAC16ΔRta | this study | a | D50.18 | F | no | FVB/NJ | 21 | no | . |
| | this study | a | D50.7 | F | no | FVB/NJ | 35 | no | . |
| | this study | a | D50.3 | F | no | FVB/NJ | 64 | **yes (4)** | . |
| | this study | a | D50.10 | F | **yes** | FVB/NJ | Euthanized (400) | no | **yes (5)** |
| | this study | b | D50.14 | M | **yes** | FVB/NJ | Euthanized (300) | no | no |
| | this study | b | D50.29 | M | no | FVB/NJ | 21 | no | . |
| | this study | b | D50.30 | M | no | FVB/NJ | 27 | no | . |
| | this study | b | D50.13 | M | **yes** | FVB/NJ | Euthanized (303) | no | no |
| | this study | b | D50.15 | F | no | FVB/NJ | 104 | **yes** | . |

1. KSHV-BAC16 was provided by J. Jung [PMID:22740391], KSHV-BAC16ΔMiR was provided by R. Renne [PMID: 26907327], KSHV-BAC16ΔRta, this paper.
2. The data for KSHV-BAC16 and KSHV-BAC16ΔMiR were previously published in [PMID: 38653242]
3. Different injection dates and oocyte donor
4. . = not available for terminal necropsy
5. Collapsed lung with pleural effusion
6. Permanent lines: FVB.tgKSHVΔRta and B6(N9).tgKSHVΔRta

| Tumor | LANA | Hemosiderin | Head and Neck | Lymphnode | Extranodal | Vasoformative spindle cells. | Cavernous spaces | Lymphangioma-like pattern | Thrombosis | Lymphocytes | Plasma cells | Salivary gland | Site | Associated Tissue(s) |
|---|---|---|---|---|---|---|---|---|---|---|---|---|---|---|
| 472 | Y | Y | Y | Y | Y | Y | Y | Y | N | Y | Y | N | Anterior | Lymph node |
| 502tu2 | Y | Y | Y | N | NA | N | Y | Y | Y | Y | Y | N | Left neck | Striated muscle |
| 506tu1 | Y | Y | Y | N | NA | Y | N | Y | NA | Y | Y | y | Middle | Salivary gland |
| 506tu2 | Y | Y | N | N | NA | Y | Y | Y | NA | Y | Y | N | Muscle | Soft tissue |
| 506tu3 | Y | N | N | Y | N | Y | N | Y | NA | Y | Y | y | Skin | Lymph node. Salivary gland. |
| 522tu1 | Y | Y | Y | N | NA | N | Y | Y | Y | Y | Y | y | Left neck | Salivary gland |
| 522tu2 | Y | Y | Y | N | NA | Y | Y | Y | Y | Y | Y | y | Left neck | Salivary gland |
| 535tu1 | Y | N | N | Y | N | Y | N | Y | N | Y | Y | N | Left flank | Lymph node |
| 541tu1 | Y | N | Y | N | NA | Y | Y | Y | N | Y | NA | N | Right neck | Soft tissue |
| 543 | Y | Y | Y | N | NA | N | Y | Y | N | Y | Y | N | Right neck | Soft tissue |
| 623LNMLN | NA | N | Y | Y | N | N | Y | Y | N | Y | N | y | Middle | Lymph node. Salivary gland. |
| 623Tu1 | NA | Y | Y | Y | Y | Y | Y | Y | N | Y | N | y | Right neck | Salivary gland |
| 630RNMLN | NA | N | Y | Y | N | N | N | Y | N | Y | Y | y | Right | Lymph node. Salivary gland. |
| 630Tu1 | NA | Y | Y | Y | Y | Y | Y | Y | N | Y | N | y | Left neck | Lymph node. Salivary gland. |
| 630Tu2 | NA | Y | Y | N | NA | Y | Y | Y | N | Y | N | N | Left neck | Soft tissue |
| 650FLTu | NA | Y | N | N | NA | Y | Y | Y | N | Y | Y | N | Flank | Soft tissue. Fat. Fibrous stroma. Ducts. Striated muscle. |
| 650HLN | NA | Y | N | N | NA | Y | N | Y | N | Y | Y | N | Next to heart Mediastinal lymph node | Soft tissue. Fat. Fibrous stroma. Striated muscle. |
| 650NTu1 | NA | Y | Y | N | NA | Y | Y | Y | N | Y | Y | N | Neck | Soft tissue. Fat. Fibrous stroma. |
| 650NTu3 | NA | Y | Y | Y | Y | N | Y | Y | N | Y | Y | y | Neck | Lymph node. Salivary gland. |
| 651LALN | NA | Y | N | Y | N | N | N | Y | N | Y | Y | N | Left axilla | Lymph node |
| 651LNR | NA | Y | Y | Y | N | N | N | Y | N | Y | Y | N | Right neck | Lymph node |
| 651smTu | NA | Y | Y | Y | Y | Y | N | Y | Y | Y | Y | N | Right neck | Lymph node |
| 690Tu1 | NA | Y | Y | Y | Y | Y | Y | Y | N | Y | Y | y | Middle neck | Lymph node. Salivary gland. Adipose. |
| 707TNTu | NA | Y | Y | Y | Y | Y | Y | Y | N | Y | Y | y | Neck | Lymph node. Adjacent salivary gland and fat. |
| 730Ntu | NA | Y | Y | N | NA | Y | Y | Y | N | Y | Y | y | Left neck | Salivary gland and fibrofat |
| 730Skin | NA | Y | N | N | NA | Y | Y | Y | N | Y | Y | N | Skin over neck tumor | Skin |
| 738Ftu | NA | Y | Y | N | NA | Y | Y | Y | Y | Y | Y | N | Face and jaw muscle | Striated muscle |
| 738RNLN | NA | Y | Y | Y | Y | Y | Y | Y | N | Y | Y | y | Right neck | Lymph node. Adjacent salivary gland and fat. |
| 764Headtu | NA | Y | Y | N | NA | Y | N | Y | N | Y | Y | N | Skin above left eye | Skin |
| 764LNTu | NA | Y | Y | Y | Y | Y | N | Y | N | Y | Y | y | Left neck connected to artery | Two lymph nodes, Adjacent salivary glands and fat. |
| 1022tu | Y | Y | Y | Y | Y | N | Y | Y | Y | Y | Y | y | Neck under chin right side | Lymph node and salivary gland. |
